# Nano3P-seq: charting the coding and non-coding transcriptome at single molecule resolution

**DOI:** 10.1101/2024.11.20.624491

**Authors:** Oguzhan Begik, Leszek P Pryszcz, Adnan Muhammad Niazi, Eivind Valen, Eva Maria Novoa

## Abstract

RNA polyadenylation is crucial for RNA maturation, stability and function, with polyA tail lengths significantly influencing mRNA translation, efficiency and decay. Here, we provide a step-by-step protocol to perform Nanopore 3’ end-capture sequencing (Nano3P-seq), a nanopore-based cDNA sequencing method to simultaneously capture RNA abundances, tail composition and tail length estimates at single-molecule resolution. Taking advantage of a template switching-based protocol, Nano3P-seq can sequence any RNA molecule from its 3’ end, regardless of its polyadenylation status, without the need for PCR amplification or RNA adapter ligation. We provide an updated Nano3P-seq protocol that is compatible with R10.4 flowcells, as well as compatible software for polyA tail length and content prediction, which we term *PolyTailor*. We demonstrate that *PolyTailor* provides accurate estimates of transcript abundances, tail lengths and content information, while capturing both coding and non-coding RNA biotypes, including mRNAs, snRNAs, and rRNAs. This method can be applied to any RNA sample of interest (e.g. poly(A)-selected, ribodepleted, total RNA), and can be completed in one day. The Nano3P-seq protocol can be performed by researchers with moderate experience in molecular biology techniques and nanopore sequencing library preparation, and basic knowledge of linux bash syntax and R programming. This protocol makes Nano3P-seq accessible and easy to implement by future users aiming to study the tail dynamics and heterogeneity of both coding and non-coding transcriptome in a comprehensive and reproducible manner.

**Key Papers:** Beğik O, Diensthuber G, Liu H, Delgado-Tejedor A, Kontur C, Niazi AM, Valen E, Giraldez AJ, Beaudoin JD, Mattick JS, Novoa EM. Nano3P-seq: transcriptome-wide analysis of gene expression and tail dynamics using end-capture nanopore cDNA sequencing. *Nature Methods* **20**, 75–85 (2023).

*https://doi.org/10.1038/s41592-022-01714-w*

Delgado-Tejedor A, Medina M, Begik O, Cozzuto L, Lopez J, Blanco B, Ponomarenko J, Novoa EM. Native RNA nanopore sequencing reveals antibiotic-induced loss of rRNA modifications in the A- and P-sites. *NatComm* **15**, 10054 (2024). *https://doi.org/10.1038/s41467-024-54368-x*

## Introduction

RNA polyadenylation is a critical process in the regulation of gene expression and RNA stability, influencing various cellular processes, including nuclear export, translational efficiency and RNA degradation ^1,2^. Recent studies have underscored the heterogeneity in poly(A) tails and their significance in fine-tuning gene expression across diverse biological contexts ^3,4^. Variations in poly(A) tail length and composition are now recognized as essential features for understanding transcriptome complexity and the regulation of gene expression at the single-molecule level.

Nanopore sequencing is a revolutionary long-read RNA sequencing technology that can capture both cDNA and native RNA molecules, facilitating the analysis of gene expression patterns and transcriptomic heterogeneity across a broad range of biological processes ^5^. This technology relies on the passing of RNA or DNA molecules through nanometer-scale pores, referred to as ‘nanopores’, resulting in detectable alterations in the electrical current signals that can be translated into nucleotide sequences through the use of machine learning algorithms ^6^. Due to its ability to capture individual full-length RNA molecules, including their tails, nanopore sequencing offers the possibility to examine poly(A) tail dynamics with unprecedented resolution.

With the discontinuation of the previous nanopore sequencing chemistry and flowcells (R9.4), PCR-free direct cDNA sequencing kits are no longer commercially offered by Oxford Nanopore Technologies (ONT). An alternative approach to characterize RNA populations using ONT consists of sequencing the native RNA molecules, known as ‘direct RNA sequencing’, which has the advantage of providing direct, real-time, and single-molecule insights into RNA abundances and even RNA modifications ^7^. However, this approach typically requires high input amounts (300 ng of polyA-selected RNA material) and is limited to sequencing polyA+ material, due to the initial ligation step that relies on oligo(dT) annealing of the DNA adapter to the RNA sample of interest. Consequently, it is unable to simultaneously capture information from both the coding and non-coding transcriptome while retaining the polyA tail length information. To overcome these limitations, we recently proposed an alternative RNA sequencing approach, which we termed Nanopore 3’ End-capture sequencing (Nano3P-seq) ^8^, which can capture both the coding and non-coding transcriptome, as well as provide accurate measurements of RNA abundances, tail lengths and tail composition and heterogeneity, with single molecule resolution, without the need of PCR amplification.

### Development of the protocol

Here we present an updated and improved version of the Nano3P-seq protocol (**Figure 1)**, with five key differences compared to the original protocol ^8^: (i) replacement of TGIRT enzyme (Ingex) with Induro RT (NEB) at reverse transcription step, due to the discontinuation of TGIRT enzyme; (ii) replacement of direct cDNA sequencing kit (DCS109) and native barcoding expansion kits (NBD104), which are both now deprecated, with the barcoding kit 24 v14 (NBD114.24), making the libraries compatible with the latest R10.4 flow cells; (iii) replacement of R9.4 flowcells, which are now deprecated, for R10.4 flowcells; (iv) modification of the reverse transcription oligonucleotides –previously terminated with a TTC sequence and occasionally misread as TTT, resulting in an overestimation of poly(T) length–, to increase the accuracy and recovery of reads with tail length estimations, (v) development of a novel software in-house tool for the analysis of polyA tail lengths and tail heterogeneity, *PolyTailor*, which is compatible with *dorado* basecalling software (replacing deprecated *guppy* basecalling software), pod5 files (instead of deprecated FAST5 files) and R10 flowcells (instead of deprecated R9.4 flowcells). We find that this updated protocol performs comparably well to the original protocol in terms of accuracy of RNA abundances, tail length estimations and recapitulation of tail length composition. Moreover, *PolyTailor* allows straightforward analysis of tail lengths and tail composition, making Nano3P-seq data analysis easy to implement by future users.

**Figure 1.**
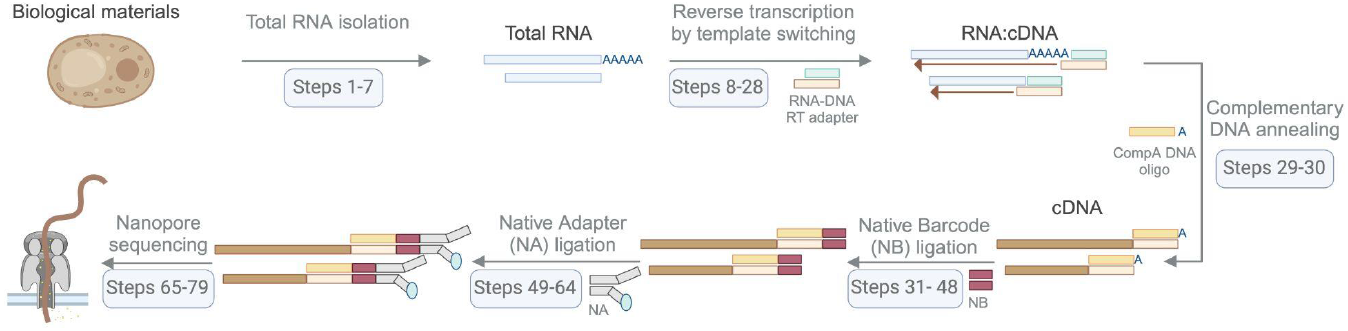
Schematic overview of the Nano3P-seq library preparation workflow. In this example, yeast total RNA is processed to prepare a Nano3P-seq library. The main steps include: (i) Total RNA isolation (Steps 1–7): extraction of total RNA from biological samples; (ii) Reverse transcription by template switching (Steps 8–28): conversion of RNA to cDNA while preserving poly(A) tail information; (iii) Complementary DNA annealing for adapter ligation (Steps 29–30); (iv) Native barcode ligation (Steps 31–48): addition of barcodes to enable sample multiplexing; (v) Native adapter ligation (Steps 49– 64): preparation of cDNA for sequencing by adding adapters; and (vi) Nanopore sequencing (Steps 65–79): real-time sequencing on a nanopore sequencing platform

### Applications of the Nano3P-seq protocol

Nano3P-seq is a simple and robust method that relies on nanopore cDNA sequencing for accurately estimating transcript levels, tail lengths and tail composition heterogeneity in individual molecules, with minimal library preparation biases, both in the coding and non-coding transcriptome. By employing a template switching-based library preparation approach, Nano3P-seq can sequence any RNA molecule from its 3’ end, regardless of its polyadenylation status, without the need for PCR amplification or ligation of RNA adapters. Notably, Nano3P-seq can capture multiple layers of transcriptomic information in one single sequencing run. More specifically, it provides highly accurate transcript abundance quantification (for both polyadenylated and non-polyadenylated RNAs simultaneously), transcriptome-wide polyA tail length estimations, and tail composition heterogeneity (e.g., internal G modifications, terminal uridylation, etc). Therefore, it can provide direct associations between transcriptomic changes and post-transcriptional modifications in the tail (**Figure 2**).

**Figure 2.**
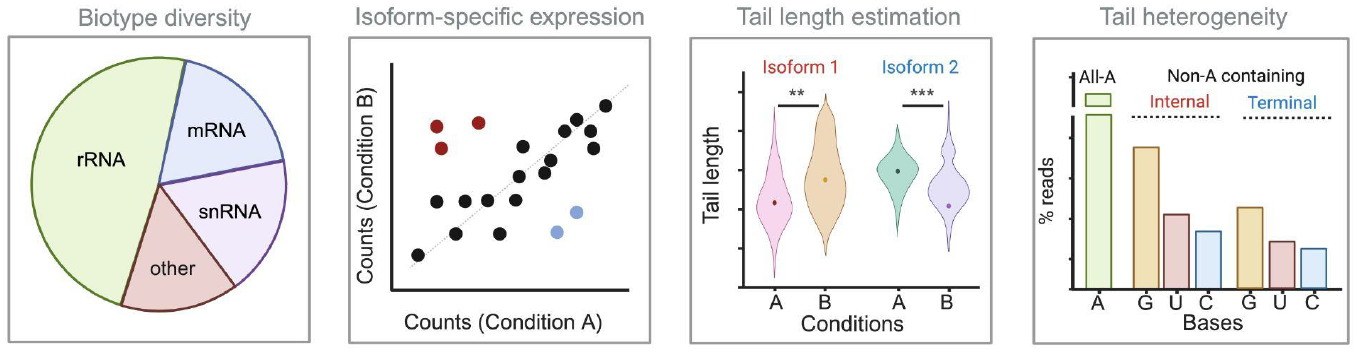
Applications of Nano3P-seq protocol. Nano3P-seq enables comprehensive analysis of both polyadenylated and non-polyadenylated RNAs, providing insights into RNA biotype diversity, isoform-specific expression levels, poly(A) tail length estimates, and tail composition heterogeneity (left to right panels). *Biotype diversity*: A pie chart representing the proportion of each RNA biotype identified in the sample, which includes mRNA, rRNA, snRNA, and other non-coding RNAs, highlighting the ability of Nano3P-seq to capture diverse RNA biotypes within a same sample and experiment. *Isoform-specific expression*: A scatter plot illustrating isoform expression levels between two conditions (Condition A and Condition B). *Tail length estimation*: Violin plots illustrating the distribution of poly(A) tail lengths for two isoforms (Isoform 1 and Isoform 2) of a given gene, under two conditions (A and B). *Tail heterogeneity*: A bar plot depicting heterogeneity in poly(A) tail composition, indicating the percentage of reads containing non-A residues (G, U, C) in internal and terminal positions.

### Comparison with other methodologies

Compared to other existing Illumina and PacBio RNA-seq approaches, Nano3P-seq offers the following advantages: (i) it will quantify a transcript even if the RT enzyme drops (e.g. due to RNA modification or RNA structure); (ii) it will not bias the RNA abundances based on the length of the RNA molecule; (iii) it can quantify both polyadenylated and non-polyadenylated transcripts; (iv) it does not require PCR amplification or second strand synthesis steps, therefore avoiding the biases known to occur associated to these steps ^9–11^; (v) the library preparation is much faster and simpler than when using orthogonal approaches, as it does not require second strand synthesis nor PCR amplification steps. In addition, compared to Illumina-based methods: (vi) it shows improved accuracy in per-gene and per-isoform quantification levels (correlation between observed and expected abundances when using synthetic standards^12^ is 0.93 with Nano3P-seq, compared to 0.89 using Illumina-based approaches^13^); (vii) it provides quantifications both at per-gene and per-isoform level; (vii) it captures polyA tail length at per-transcript (isoform) resolution, whereas Illumina-based methods can only achieve per-gene resolution; (viii) it captures tail composition heterogeneity with high accuracy. A major limitation of Nano3P-seq, however, is that it requires 50 ng of input material to start the library, as there is no PCR amplification of the material.

Finally, we should note that while earlier PacBio-based methods, such as FLAM-seq^14^ or PAIso-seq^15^, were unable to identify 3′ terminal modifications, more recent protocols, including PAIso-seq2 ^16^, FLEP-seq ^17^ and FLEP-seq2 ^18^ have overcome this limitation. Similarly, tail heterogeneity can now be characterized in ONT direct RNA sequencing datasets using the toolkit *Ninetails* ^*19*^.

### Experimental design

#### Starting material

Nano3P-seq can be adapted to any RNA input due to its ability to capture RNAs regardless of their 3’ end sequence context. Therefore, samples such as total RNA, ribodepleted RNA, poly(A)-selected RNA, *in vitro* transcribed RNAs or other forms of synthetic RNAs can be subjected to Nano3P-seq. In this work, we have used total RNA isolated from *S. cerevisiae*, as well as synthetic DNA standards with known tail lengths and variations in their tail composition^13^, to assess both the accuracy in tail length and tail composition predictions.

We should note that additional adjustments are required if the purpose of using Nano3P-seq is to capture transfer RNAs (tRNAs) or other highly structured and/or modified RNAs, as these are typically not efficiently reverse transcribed with common reverse transcriptases and/or buffers (see **Reaction buffers** section). Furthermore, an adjusted amount of beads required for cleaning up the library might be needed when the input material consists of small RNAs (see **Bead cleanup and size selection** section).

#### Template switching

Template switching is a reverse transcription (RT) mechanism by which group II intron-encoded reverse transcriptases, such as TGIRT ^8,20^ (Ingex) or Induro RT (NEB, cat. no. M0681S), are able to initiate RT via an RNA-DNA heteroduplex adapter with a single degenerate nucleotide overhang ^21^. Thus, template switching allows reverse transcribing RNA molecules regardless of their 3’ end sequence. We should note that the original Nano3P-seq protocol relied on TGIRT for template-switching reaction, however with the commercial discontinuation of this enzyme, the protocol has been updated to use Induro RT enzyme. We should note that the use of template switching can bias the biotype proportion present in the sample ^22,23^ depending on the enzyme and buffer of choice, among other features.

#### Reaction buffers

The buffer used in Nano3P-seq RT step is a commercial buffer (Induro Buffer) to reverse transcribe most of the RNA biotypes, except for highly structured and modified RNAs, such as tRNAs. We should note that previous studies using TGIRT have demonstrated that the use of “low salt buffer” during RT enhances the reverse transcription of tRNA populations ^24,25^.

#### Bead cleanup and size selection

After reverse-transcription and RNA digestion, cDNA products are cleaned using Ampure XP beads (Agencourt). At this point, size selection can be performed using a specific amount of beads. If the aim is to enrich long RNAs, a lower amount of beads should be used.

#### Library preparation

To ligate native barcodes to each sample, a complementary DNA (CompA_DNA, see **Oligonucleotides** section below) with an adenine overhang is annealed to the purified cDNAs, enabling the ligation of native barcodes to cDNAs with Blunt/TA ligase (NEB). At this step, the user should ensure not to exceed the recommended cDNA amount per barcode, which is 200 fmol (see **Step 31**). It is also important to note that multiplexing a higher number of libraries per flowcell will lead to a decreased number of reads per sample.

Hence, it is advised to multiplex fewer libraries if a higher amount of reads per each sample is desired. Once the barcoding step is inactivated with EDTA, libraries are pooled together, followed by another cleanup with Ampure XP beads. The final step of library preparation involves the ligation of native adapters to the pool of barcoded samples.

Of note, while this protocol is designed for multiplexed libraries, barcoding steps can be skipped if desired, and cDNA products can be directly ligated to sequencing adapters. In this case, we recommend users to employ the Ligation Sequencing Kit v14 (ONT, SQK-LSK114), which offers a sequencing adapter compatible with our protocol.

#### Nanopore sequencing

Once eluted, the library is kept on ice until loaded onto a flow cell. Flowcells must be carefully primed without introducing air into the priming port, which would otherwise lead to the formation of bubbles in the flowcell, causing a decrease in the number of pores that are available for sequencing, and consequently, sequencing yield. The duration of the nanopore sequencing runtime can vary depending on library and flowcell quality. A Nano3P-seq run takes a maximum of 72 hours to reach saturation. Analysis is performed once the sequencing run is finished and all data has been saved.

### Data analysis

Nano3P-seq data analysis involves the use of several softwares, some of them being common to any nanopore sequencing run (**Figure 3**). First, raw nanopore data (POD5 files) is basecalled (and demultiplexed) using *Dorado* basecaller, which generates SAM/BAM files that contain necessary information to predict polyA tail lengths (move table). Then, basecalled reads are aligned to the reference genome using *minimap2* transferring move tables in the process (the alignment has to be performed separately, because *dorado* basecaller does not support splice-aware alignment to date). Finally, *polyTailor*, a tool we developed *in-house* for Nano3P-seq data analysis, is used to estimate polyA tail length and tail content for every read, overlap with known/predicted TES and read-to-transcript correspondence.

**Figure 3.**
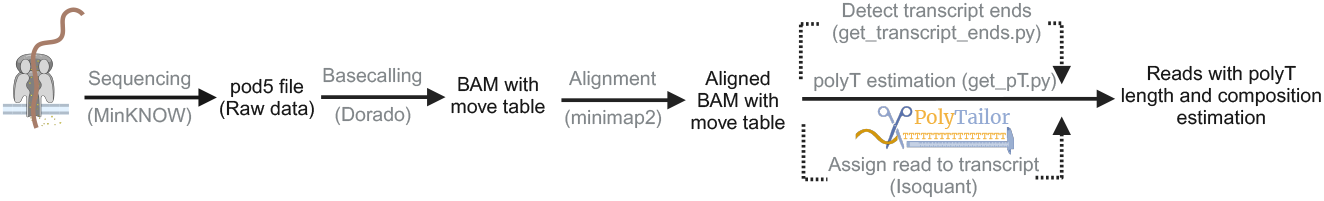
Schematic flowchart of Nano3P-seq data analysis steps. This diagram illustrates the data processing pipeline for Nano3P-seq datasets, to extract poly(A) tail length and composition information. Raw nanopore sequencing data (pod5 files) are basecalled using *dorado*, generating BAM files with move (mv) tables as output. The BAM files are then aligned to the reference transcriptome using minimap2, resulting in aligned BAM files with move tables, which facilitates subsequent transcript end detection and poly(A) tail analysis. We then use *PolyTailor* to extract information from the aligned BAM files, using different scripts: i) *get_transcript_ends*.*py* detects transcript ends; ii) *get_pT*.*py* estimates poly(A) tail length and composition. Finally, we use *Isoquant* to determine read-to-transcript associations. Using these 3 outputs, PolyTailor then generates a final file that contains readID, transcript ID, poly(A) tail length and tail heterogeneity information.

We should note that Nano3P-seq protocol also captures fragmented RNAs, which may confound the polyA tail length analysis predicting reads arising from fragmented transcripts as reads without polyA tails. Therefore, we recommend narrowing the analysis only to those reads that can be associated with known or predicted transcript end sites (TES). Novel TES can be predicted using *get_transcript_ends*.*py* from the PolyTailor tool. Script. The user may further narrow the analysis to those reads that match expected exon-intron structures using the software *IsoQuant* ^*26*^. Finally, *get_pT*.*py from the PolyTailor tool* is used to obtain tail predictions.

### Expertise needed to implement the protocol

The Nano3P-seq protocol requires moderate experience with molecular biology experimental techniques. Experience with nanopore sequencing library preparation and sequencing is not essential but is highly recommended. RNA handling requires special attention to avoid degradation due to RNase contamination. Furthermore, users should have access to a high-performance computing cluster or cloud services to perform computation intensive tasks. Basic knowledge of linux bash computing and R programming to be able to run the software used for data analysis is also required.

### Limitations

The Nano3P-seq protocol uses a template switching mechanism to initiate reverse transcription from any RNA molecule with a free 3’end. Thus, Nano3P-seq also can capture some fragmentation products arising from RNA being exposed to high temperatures cleaved by RNases. Fragmented transcripts can be filtered out by the user by narrowing the analysis to reads matching known TES and/or expected exon-intron structures. Notably, *polyTailor* software allows the user to identify novel TES and transcripts from the data that is being analyzed, and is applicable to organisms lacking genome annotations.

Finally, we should note that it is experimentally challenging to prepare a library that efficiently captures both very short RNAs, such as transfer RNAs (tRNAs), and long RNAs at the same time, due to the limitations of bead cleanup and ligation steps. However, the user can choose to employ low salt RT buffers ^22,27^ and high amounts of beads during the cleanup steps to enhance the recovery of small and highly modified RNAs, such as tRNAs.

## Materials

### Biological Materials

#### Yeast culturing and total RNA extraction

*Saccharomyces cerevisiae* (strain BY4741) was grown at 30ºC in standard YPD medium (1% yeast extract, 2% Bacto Peptone and 2% dextrose, all wt/vol).

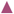 **CRITICAL** Our procedure is suitable for any material RNA can be extracted from. However, users may need to adjust the amount of starting material based on the average yield of RNA.

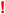 **CAUTION** Animal experiments should be performed according to the relevant institutional and national guidelines and regulations.

#### Reagents

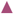 **CRITICAL** It is essential to use the library preparation kit offered by Oxford Nanopore Technologies (ONT) in order to perform this protocol.

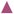 **CRITICAL** An ONT account is needed to order ONT products

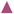 **CRITICAL** This protocol is suitable for barcoded samples.

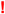 **CAUTION** TRIzol and chloroform are harmful reagents and must be handled carefully. Please follow the guidelines to handle these reagents and dispose them properly.

- RNaseZap™ RNase Decontamination Solution (ThermoFisher, cat. no. AM9780)
- TRIzol Reagent (Invitrogen, cat. no. 15596018)
- Acid washed and autoclaved glass beads 425-600 µm (Sigma, cat.no. G8772)
- Chloroform (Invitrogen, cat. no. 15596018)
- Ethanol, pure 200 proof, for molecular biology (Sigma-Aldrich, cat. no. E7023-500mL)
- Sodium Acetate 3M, pH 5.2 (ThermoFisher, cat.no. R1181)
- Turbo DNase (ThermoFisher, cat.no. AM2238)
- 1 M Nuclease-free Tris-Cl pH 7.5 (ThermoFisher, cat. no. 15567027)
- 5 M Nuclease-free NaCl (ThermoFisher, cat. no. AM9760G)
- Nuclease-free water (ThermoFisher, cat. no. AM9922)
- Induro (NEB, cat. no. M0681S)
- RNase Inhibitor, Murine (NEB, cat. no. M0314L)
- 10 mM dNTP solution (ThermoFisher, cat. no. R0191)
- RNase Cocktail Enzyme Mix (ThermoFisher, cat. no. AM2286)
- AMPure XP Reagent (Agencourt, cat. no. A63881)
- Native Barcoding Kit 24 V14 (ONT, cat. no. SQK-NBD114.24)
- Blunt/TA Ligase Master Mix (NEB, cat. no. M0367)
- NEBNext Quick Ligation Module (NEB, cat. no. E6056)
- BSA (ThermoFisher, cat. no. AM2616)

#### Equipment and Materials

- 0.2 ml thin-walled PCR tubes (Starlab, cat. no. A1402-3700)
- 1.5 mL DNA LoBind Tubes (Eppendorf, cat. no. 0030108051)
- 10-, 100-, 200- and 1,000 μl filter barrier tips pipette tip, filter, sterile (Axygen, mod. nos. TF-300-R-S, TF-100-R-S, TF-200-R-S, TF-1000-L-R-S)
- TapeStation (Agilent, mod. no. G2991BA) and accompanying reagents: RNA Screen-Tape Sample Buffer, RNA ScreenTape Ladder, RNA Screen-Tape, optical tube strip caps (8” strip), optical tube strips (8” strip), and loading tips.
- Qubit 3.0 fluorometer (Thermo Fisher Scientific, mod. no. Q33216) or Qubit 4.0 fluorometer, Thermo Fisher Scientific, mod. no. Q33238)
- Nanodrop 2000 (Thermo Fisher Scientific, mod. no. ND2000)
- Eppendorf ThermoMixer C with Thermo top (Eppendorf, mod. no. 5382000023)
- Tube Rotator (MACSmix, mod. no. 130-090-753)
- DynaMag-2 magnet for 1.5 ml microtube (Thermo Fisher Scientific, mod. no. 12321D)
- Refrigerated centrifuge (e.g., Eppendorf, mod. no. 5430R)
- Bench top centrifuge (e.g., Eppendorf, mod. no. minispin)
- Thermocycler (e.g., BioRad, mod. no. T100)
- Minicentrifuge (e.g., Thermo Fisher Scientific, mod. no. The Fisherbrand Mini-Centrifuge)
- Nanopore sequencing device (Oxford Nanopore Technologies, MinION/GridION/PromethION)
- Nanopore R10.4.1 flow cell (FLO-MIN114)
- An UNIX/Linux workstation with 16 GB memory and preferably GPU compatible with *dorado* basecaller

#### Oligonucleotides

- D_DNA: /5Phos/ACTTGCCTGTCGCTCTATCTGCAGAGCAGAGN (Order 100 nmol of DNA oligo with standard desalting. Dissolve the lyophilized oligo in nuclease-free water to a final concentration of 100 uM. The dissolved oligo can be stored at -20 °C for long term storage.)
- R_RNA: rCrUrCrUrGrCrUrCrUrGrCrArGrArUrArGrArGrCrGrArCrArGrGrCrArArGrU/3SpC3/ (Order this oligo with RNase-free HPLC purification method. Dissolve the lyophilized oligo in nuclease-free water to a final concentration of 100 uM. The dissolved oligo can be stored at -80 °C for long term storage.)
- CompA_DNA: CTCTGCTCTGCAGATAGAGCGACAGGCAAGTA (Order 100 nmol of DNA oligo with standard desalting. Dissolve the lyophilized oligo in nuclease-free water to a final concentration of 100 uM. The dissolved oligo can be stored at -20 °C for long term storage.)

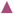 **CRITICAL** Nano3P-seq runs performed with the older chemistry (R9) use different R_RNA and D_DNA sequences. We altered the sequences to increase the efficiency of tail predictions. Deprecated primer sequences, used with R9 chemistry, can be found on the PolyTailor github page (https://github.com/novoalab/polyTailor)

#### Software

- MinKNOW (version v24.02 or later): https://community.nanoporetech.com/downloads
- Dorado (version v0.7.2 or later) : https://github.com/nanoporetech/dorado 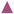 **CRITICAL** Nano3P-seq analysis pipeline has been tested on reads basecalled with Dorado v0.7.2 and more recent versions are also likely to work as expected. The latest version of Dorado is always recommended considering the base calling accuracy is continuously improving in the recent Dorado releases.
- PolyTailor (version 1.0): https://github.com/novoalab/polyTailor
- Isoquant (version 3.5): https://github.com/ablab/IsoQuant
- Samtools (version 1.19): https://github.com/samtools/htslib
- Minimap2 (version 2.28): https://github.com/lh3/minimap2

### Equipment setup

#### Example datasets

For yeast total RNA samples, download the example Nano3P-seq dataset generated from the public repository: https://public-docs.crg.es/enovoa/public/lpryszcz/src/polyTailor/test

#### Software installation

The Nano3P-seq pipeline requires a UNIX/Linux environment. To install and configure the necessary tools, please follow the instructions in **Box 1**.

##### Box 1

**Software setup for Nano3P-seq analysis** 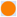 **Timing** 1 h

**PolyTailor installation**

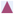 **CRITICAL** The PolyTailor software requires a UNIX/Linux environment. Users must have access to a high-performance computing cluster or Linux workstation. Users with access to Nvidia GPU / Apple silicon GPU (M1, M2, M3) will benefit from hardware-accelerated basecalling. PolyTailor requires python version 3.8 or higher.

Step 1. Install required packages

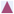 **CRITICAL** Installation instruction for *conda* can be found at https://docs.anaconda.com/miniconda/

~~~
conda create -c conda-forge -c bioconda -n polyTailor python=3.10 samtools
minimap2 isoquant
conda activate polyTailor
pip install matplotlib numpy parasail pybedtools pysam pandas scipy seaborn
htseq
~~~

Step 2. Download PolyTailor from github

~~~
mkdir -p ∼/src && cd ∼/src
git clone https://github.com/novoalab/polyTailor.git
~~~

Following the steps above, PolyTailor will be ready to use. Please refer to the GitHub repository in case you encounter any issues when running the software.

**Installation of other ONT softwares required for Nano3P-seq data analysis**

These tools can be installed following ONT’s instructions.

Download and install MinKNOW from the following link: https://community.nanoporetech.com/downloads

Download and install Dorado from the following link: https://community.nanoporetech.com/downloads

For Linux x64 system, you can follow these instructions:

~~~
mkdir -p ∼/src && cd ∼/src
wget https://cdn.oxfordnanoportal.com/software/analysis/dorado-0.7.2-linux-x64.tar.gz
tar xpfz dorado-0.7.2-linux-x64.tar.gz
echo 'export PATH=∼/src/dorado-0.7.2-linux-x64/bin:$PATH’ >> ∼/.bashrc
source ∼/.bashrc
dorado download --directory ∼/src/dorado/models --model dna_r9.4.1_e8_hac@v3.3
dorado download --directory ∼/src/dorado/models --model dna_r9.4.1_e8_sup@v3.3
dorado download             --directory       ∼/src/dorado/models       --model
dna_r10.4.1_e8.2_400bps_hac@v5.0.0
dorado download             --directory       ∼/src/dorado/models       --model
dna_r10.4.1_e8.2_400bps_sup@v5.0.0
~~~

### Procedure

#### Total RNA extraction 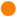 Timing 3 h

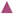 **CRITICAL** We have used *Saccharomyces cerevisiae* in this publication. However, this method works for many types of starting materials.

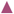 **CRITICAL** RNA molecules are prone to digestion by ribonucleases. In order to prevent RNA samples from being degraded, all reagents must be purchased or prepared nuclease free. Before the procedure, the workbench must be treated with RNaseZap (ThermoFisher) solution. Gloves must be changed frequently throughout the procedure to avoid RNase contamination. RNA samples and enzymes must be kept on ice if needed. RNA extraction using TRIzol in combination with RNeasy MinElute kit is recommended, however other methods can also be used for RNA extraction.

1. Resuspend the flash frozen *Saccharomyces cerevisiae* (strain BY4741) cells grown overnight at 30ºC in 700 µL TRIzol (Life Technologies) with 350 µL acid washed and autoclaved glass beads (425-600 µm, Sigma). Disrupt the cells using a vortex on top speed for 7 cycles of 15 seconds (Chill the samples on ice for 30 seconds between cycles).
2. Afterwards, incubate the samples at room temperature for 5 minutes and add 200 µL chloroform. After briefly vortexing the suspension, incubate the samples for 5 minutes at room temperature. Then centrifuge at 14,000 g for 15 minutes at 4ºC and transfer the upper aqueous phase to a new tube.
3. Precipitate RNA with 2X volume Molecular Grade Absolute ethanol and 0.1X volume Sodium Acetate (3M, pH 5.2). Incubate the samples for 1 hour at -20ºC and centrifuge at 14,000 g for 15 minutes at 4ºC.
4. Wash the pellet with 70% ethanol (vol/vol) and resuspend it with nuclease-free water after air drying for 5 minutes on the benchtop.
5. Measure the total RNA with the NanoDrop 2000 Spectrophotometer.
6. Treat the total RNA with Turbo DNase (Thermo) (2 ul enzyme for 50 ul reaction of 200 ng/ul RNA) at 37ºC for 15 minutes, with a subsequent RNAClean XP bead cleanup (Following manufacturer’s recommendations).
7. Check the size distribution of the extracted RNA using a TapeStation (Agilent). A high-quality total RNA should have an RNA Quality Number (RIN) higher than 8.5. 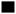 **PAUSE POINT** Extracted total RNA can at -80 ° C for long term.

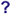 **TROUBLESHOOTING**

#### Nano3P-seq library preparation

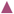 **CRITICAL** RNA molecules are prone to digestion by ribonucleases. In order to prevent RNA samples from being degraded, all reagents must be purchased or prepared as RNase-free. Before starting the library preparation, the workbench must be treated with RNaseZap (ThermoFisher) solution. Gloves must be changed frequently throughout the procedure to avoid RNase contamination. RNA samples and enzymes must be kept on ice if needed. RNase Inhibitors should be added to each reaction in order to inactivate RNases.

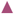 **CRITICAL** For each bead cleanup step, transfer the supernatant into a new clean eppendorf tube and keep on ice once bead suspension incubated with cDNAs are placed on the magnetic stand. In case of problems related to bead cleanup, the supernatant can be used to troubleshoot and recover any unbound cDNAs (see **Troubleshooting Table**).

#### Reverse Transcription by template switching 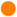 Timing 2.5 h

8. Add the following reaction components in a 0.2 mL PCR tube in the order specified: 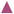 **CRITICAL STEP** To initiate reverse transcription by template switching, we must first pre-anneal the R_RNA and D_DNA oligos.

| Reagent | Volume (ul) | Final Concentration |
| --- | --- | --- |
| Tris-Cl pH 7.5 (0.1 M) | 1 | 0.01 M |
| NaCl (0.5 M) | 1 | 0.05 M |
| RNase Inhibitor, Murine (40 U/ul) | 0.5 | 2 U/ul |
| R_RNA (10 uM) | 1 | 1 uM |
| D_DNA (10 uM) | 1 | 1 uM |
| Nuclease-free water | 5.5 | - |
| Total volume | 10 | - |

9. Heat the mixture to 82°C for 2 mins and ramp down to 25°C at 0.1°C/s in the thermal cycler. 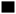 **PAUSE POINT** Pre-annealed oligos can be stored at-80 ° C for long term
10. Mix RNA sample and pre-annealed oligos in a 0.2 mL PCR tube 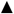**CRITICAL** Prepare one reaction for each sample to be barcoded.

| Reagent | Volume (ul) | Final Concentration |
| --- | --- | --- |
| Total RNA | Up to 50 ng | - |
| Pre-annealed oligos (1 uM) | 2 | 0.1 uM |
| 5X Induro RT Reaction Buffer | 4 | 1X |
| RNase Inhibitor, Murine (40U/ul) | 1 | 2 U/ul |
| Induro Reverse Transcription (200 U/ul) | 1 | 10 U/ul |
| Nuclease-free water | Up to 19 |  |
| Total | 19 |  |

11. Incubate the reaction at room temperature for 30 minutes before proceeding to the next step. Add 1 ul dNTP (10 mM) and proceed to the next step.
12. Incubate the reaction at 55°C for 30 minutes for reverse-transcription and 95°C for 1 minute for inactivation. In the meantime, equilibrate the Ampure XP beads at least 30 minutes before use.
13. Move reaction to ice. Add 19 ul water.
14. Add 1 ul RNase Cocktail Enzyme Mix to each tube and incubate the reaction at 37°C for 10 minutes.
15. Move reaction to ice and proceed to the clean-up step.
16. Add 32 ul of Ampure XP beads (0.8 x volumes) to each reaction. Mix by flicking. 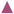 **CRITICAL STEP** The bead volume depends on the expected size of your library. Please refer to the size selection chart of the manufacturer.
17. Incubate 10 minutes at room temperature.
18. Spin down the tube and place it on the magnetic stand until the solution is clear (∼ 2 minutes).
19. Keep the tube on the magnetic stand. Remove the supernatant carefully without disturbing the beads.
20. Add 200 ul freshly prepared 70% ethanol (vol/vol) to the tube. Wait for 30 seconds and discard all the supernatant. 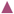 **CRITICAL STEP** Beads should be always kept on the magnetic stand during washing steps and they should not be resuspended.
21. Repeat the previous step.
22. Remove the ethanol completely by spinning down the tube, placing it back on the magnetic stand and removing the excess liquid.
23. Air-dry the beads on the magnetic stand for 30 seconds. 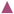 **CRITICAL STEP** Beads should not be over-dried since they will crack and it will decrease the cDNA recovery.
24. Resuspend beads with 16 ul nuclease-free water.
25. Incubate at room temperature for 10 minutes.
26. Place the tube on the magnetic rack and wait until the solution is clear.
27. Transfer the supernatant into a new 0.2 mL tube.
28. Take 1 ul of the elute and quantify using Qubit fluorometer with DNA HS Assay. The expected yield for each sample in total should be 5-10 ng from 50 ng starting material. 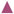 **CRITICAL STEP** Reverse transcription product size can be assessed using TapeStation RNA Assay (Agilent). Although size estimation will not be accurate, it will give a rough idea to assess the number of cDNA molecules and the efficiency of the reverse transcription reaction. 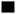 **PAUSE POINT** Reverse transcription product can be stored at -20 °C for long term.

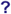 **TROUBLESHOOTING**

#### Complementary DNA annealing and Native Barcode Ligation 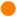 Timing 1 hour

29. Add the following reaction components in a 0.2 mL tube in the order specified and mix by flicking. 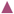 **CRITICAL STEP** This step is essential to have a double-stranded DNA with an A overhang on the 5’ end of each reverse-transcription product, which will aid for the ligation of the adapter.

| Reagent | Volume (ul) | Final Concentration |
| --- | --- | --- |
| cDNA | 15 | - |
| Tris-Cl pH 7.5 (0.1 M) | 2.25 | 0.01 M |
| NaCl (0.5 M) | 2.25 | 0.05 M |
| CompA_DNA (10 uM) | 1 | 0.44 uM |
| Water | 2 | - |
| Total | 22.5 | - |

30. Heat the mixture to 82°C for 2 mins and ramp down to 25°C at 0.1°C/s in the thermal cycler.
31. Add the following to the reaction tube.

30. Heat the mixture to 82°C for 2 mins and ramp down to 25°C at 0.1°C/s in the thermal cycler. 31. Add the following to the reaction tube.
| Reagent | Volume (ul) |
| --- | --- |
| cDNA - CompA_DNA | 22.5 |
| Native Barcode (NB01-24, ONT) | 2.5 |
| Blunt/TA Ligase Master Mix | 25 |
| Total | 50 |

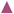 **CRITICAL STEP** It is recommended not to exceed the maximum cDNA amount of 200 fmol per barcode for the ideal ligation reaction.

32. Mix by flicking or gently pipetting and quickly spin down.
33. Incubate at room temperature for 20 minutes. In the meantime, equilibrate the Ampure XP beads at least 30 minutes before use.
34. Add 5 ul of EDTA (Provided in the ONT kit) to each tube and mix thoroughly by pipetting and quickly spin down to inactive the reaction.
35. Pool all the barcoded samples in a 1.5 mL DNA LoBind tube.
36. Mix the pooled samples with 0.5 x volume of Ampure XP beads and mix by flicking. 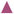 **CRITICAL STEP** The bead volume depends on the expected size of your library. Please refer to the size selection chart of the manufacturer.
37. Incubate 10 minutes at room temperature.
38. Spin down the tube and place it on the magnetic stand until the solution is clear (∼ 2 minutes).
39. Keep the tube on the magnetic stand. Remove the supernatant carefully without disturbing the beads.
40. Add 700 ul freshly prepared 70% ethanol (vol/vol) to the tube. Wait for 30 seconds and discard all the supernatant. 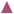 **CRITICAL STEP** Beads should be always kept on the magnetic stand during washing steps and they should not be resuspended.
41. Repeat the previous step.
42. Remove the ethanol completely by spinning down the tube, placing it back on the magnetic stand and removing the excess liquid.
43. Air-dry the beads on the magnetic stand for 30 seconds. 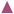 **CRITICAL STEP** Beads should not be over-dried since they will crack and it will decrease the cDNA recovery.
44. Resuspend beads with 31 ul nuclease-free water.
45. Incubate at room temperature for 10 minutes.
46. Place the tube on the magnetic rack and wait until the solution is clear.
47. Transfer the supernatant into a new 1.5 mL DNA LoBind tube.
48. Take 1 ul of the elute and quantify using Qubit fluorometer with DNA HS Assay. It will likely be below detection limit. 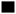 **PAUSE POINT** Pooled DNA library can be stored at -20°C for long term.

#### Native Adapter (NA) Ligation 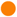 Timing 1 hour

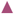 **CRITICAL** Before starting this step, thaw Long or Short Fragment buffer (LFB, SFB) and Elution Buffer (EB) at room temperature, mix by vortexing, spin down and place on ice until used. Check for any precipitation.

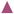 **CRITICAL** Use LFB to enrich DNA fragments of 3 kb or longer. To retain DNA fragments of all sizes, use SFB.

49. Mix the following reagents in the following order in a 1.5 mL DNA LoBind tube:

| Reagent | Volume (ul) |
| --- | --- |
| Pooled barcoded sample (Up to 200 fmol) | 30 |
| Native Adapter (NA) | 5 |
| NEBNext Quick Ligation Reaction Buffer (5X) | 10 |
| Quick T4 DNA Ligase | 5 |
| Total volume | 50 |

50. Mix gently by flicking the tube, and spin down. Incubate at room temperature for 20 minutes, In the meantime, equilibrate the Ampure XP beads at least 30 minutes before use.
51. Mix the library with 0.5 x volume of Ampure XP beads and mix by flicking. 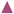 **CRITICAL STEP** The bead volume depends on the expected size of your library. Please refer to the size selection chart of the manufacturer.
52. Incubate 10 minutes at room temperature.
53. Spin down the tube and place it on the magnetic stand until the solution is clear (∼ 2 minutes).
54. Keep the tube on the magnetic stand. Remove the supernatant carefully without disturbing the beads.
55. Add 125 ul SFB/LFB to the tube. Close the tube lid, remove the tube from the magnetic stand and resuspend the beads by flicking. Return the tube to the magnetic rack and allow beads to pellet and remove the supernatant.
56. Repeat the previous step.
57. Remove the SFB/LFB completely by spinning down the tube, placing it back on the magnetic stand and removing the excess liquid.
58. Air-dry the beads on the magnetic stand for 30 seconds. 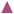 **CRITICAL STEP** Beads should not be over-dried since they will crack and it will decrease the cDNA recovery.
59. Resuspend beads with 15 ul Elution Buffer (EB).
60. Incubate at room temperature for 5-10 minutes.
61. Place the tube on the magnetic rack and wait until the solution is clear.
62. Transfer the supernatant into a new 1.5 mL DNA LoBind tube.
63. Take 1 ul of the elute and quantify using Qubit fluorometer with DNA HS Assay.
64. Make the library up to 12 uL at 10-20 fmol. 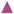 **CRITICAL STEP** Library size can be assessed using TapeStation RNA Assay. Although the size is not accurate, it will give a rough idea to assess the number of molecules in the library. Loading more than 20 fmol of DNA can reduce the rate of duplex read capture. 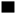 **PAUSE POINT** We recommend storing the library at 4 °C for short-term storage or repeated use and at -80°C for long term.

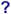 **TROUBLESHOOTING**

#### Library loading and sequencing 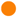 Timing 30 minutes for library loading and 48 hours for sequencing

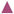 **CRITICAL** This protocol is written for minION flow cells. Necessary adjustments need to be made if the library will be loaded onto a promethION flow cell.

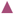 **CRITICAL** This library is only compatible with R10.4.1 flow cells (FLO-MIN114)

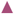 **CRITICAL** Before starting this step, thaw Sequencing Buffer (SB), Library Beads (LIB) or Library Solution (LIS, if using), Flow Cell Tether (FCT) and one tube of Flow Cell Flush (FCF) at room temperature, mix by vortexing, spin down and place on ice until used. Check for any precipitation.

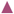 **CRITICAL** For optimal sequencing performance and improved output on FLO-MIN11 flow cells, we recommend adding Bovine Serum Albumin (BSA) to the flow cell priming mix at a final concentration of 0.2 mg/ml.

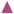 **CRITICAL** Mix the contents of LIB with a large volume of pipette to make it homogeneous just before taking.

65. To prepare the flow cell priming mix with BSA, add 5 uL BSA (50 mg/ml) and 30 uL FCT directly to a new FCF tube (1170 uL volume) and mix by pipetting.
66. Prepare the following in a 1.5 mL DNA LoBind Tube and keep on ice until loading.

| Reagent | Volume (µl) |
| --- | --- |
| Sequencing Buffer (SB) | 37.5 |
| Library Beads (LIB) | 25.5 |
| DNA Library (maximum 10-20 fmol) | 12 |
| Total | 75 |

67. Take the flow cell out of the fridge and connect it to the MinION.
68. Check the flow cell (QC it to see how many active pores there are).

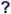 **TROUBLESHOOTING**

69. To remove any bubbles from the priming port, open the lid of the nanopore sequencing device and slide the flow cell’s priming port cover clockwise so that the priming port is visible.
70. Set a P1000 pipette to 200 uL.
71. Insert the tip into the priming port.
72. Turn the wheel until the dial shows 220-230 μl, or until you can see a small volume of buffer entering the pipette tip.
73. Load 800 uL of flow cell priming mix slowly. 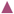 **CRITICAL STEP** Make sure not to introduce any air as it may affect the run.
74. Wait for 5 minutes and open the sample port cover.
75. Load 200 ul of priming mix more into the priming port, observing the bubbles coming out of the sample port.
76. Mix the prepared library gently by pipetting up and down just prior to loading.
77. Add 75 μl of sample to the flow cell via the SpotON sample port in a dropwise fashion. Ensure each drop flows into the port before adding the next.
78. Gently replace the SpotON sample port cover, making sure the bung enters the SpotON port, close the priming port and replace the MinION lid.
79. Start the sequencing run and keep it running for 48 hours.

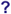 **TROUBLESHOOTING**

#### Data analysis 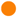 Timing 2h (hac) - 15h (sup)

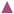 **CRITICAL** Here we use an example dataset of yeast total RNA to illustrate the data analysis steps. Related sequencing data can be accessed through https://public-docs.crg.es/enovoa/public/lpryszcz/src/polyTailor/test and annotation files can be found in the github page of PolyTailor.

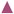 **CRITICAL** Estimated computing time required to analyse the data will vary depending on the sequencing yield, and the characteristics of the computing infrastructure available. Timings above reflect the time required to analyse 5 million reads sequenced in a MinION device in a Linux workstation with 6 CPU cores, 16 GB memory and equipped with dedicated GPU (Geforce RTX 3080 Ti).

80. Download the sequencing data and respective reference files

~~~
mkdir ∼/src/polyTailor/test; cd ∼/src/polyTailor/test
wget
~~~

https://public-docs.crg.es/enovoa/public/lpryszcz/src/polyTailor/test/

~~~
                                                  {ref,reads} -q --
show-progress -r -c -nc -np -nH --cut-dirs=6 --reject="index.html*”
~~~

81. Basecall the reads –and demultiplex if needed– by saving the *mv* table in BAM file using dorado basecaller, using either the sup or hac model.

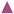 **CRITICAL** For the most accurate polyT composition calling we recommend using the latest sup model. If barcoding *--kit-name* is provided, barcodes will be reported in the barcode column. This step will benefit from hardware-accelerated basecalling for the Nvidia GPU using *-x cuda:all* (or Apple silicon M1, M2, M3).

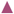 **CRITICAL** Basecalling using SUP model takes significantly longer time than HAC model (3 times longer for R9, 24 times longer for R10).

For an R9 run:

~~~
cd ∼/src/polyTailor/test
mkdir -p dorado/yeast
dorado basecaller \
   ∼/src/dorado/models/dna_r9.4.1_e8_hac@v3.3 \
   reads/yeast/R9 -r \
   --kit-name EXP-NBD104 \
   --no-trim \
   --emit-moves \
   -x cuda:all > dorado/yeast/R9.bam
~~~

For an R10 run:

~~~
dorado basecaller \
   ∼/src/dorado/models/dna_r10.4.1_e8.2_400bps_hac@v5.0.0 \
   reads/yeast/R10 -r \
   --kit-name SQK-NBD114-24 \
   --no-trim \
   --emit-moves \
   -x cuda:all > dorado/yeast/R10.bam
~~~

82. Align the reads to the genome passing dorado tags to the resulting BAM file. 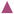 **CRITICAL** For yeast, we should reduce the intron size to 2kb using *-G2k*. For higher eukaryotes with larger genomes, this parameter should be skipped.

~~~
conda activate polyTailor
mkdir -p minimap2/yeast
for f in dorado/yeast/*.bam; do echo `date` $f;
   samtools fastq -T mv,ts,BC $f \
   | minimap2 -y -ax splice:hq -G2k ref/yeast.fa.gz - \
   | samtools sort --write-index -o minimap2/yeast/$(basename $f);
done
~~~

83. Annotate alternative transcript ends. 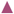 **CRITICAL** Official annotation for *S. cerevisiae* doesn’t include UTRs, therefore we extend each gene by 1000 bases using *-e 1000*. This can be skipped for genomes with complete UTR annotation.

~~~
python ../src/get_transcript_ends.py --firststrand -q 0 -e 1000 \
 -a ref/yeast.gtf.gz -b minimap2/yeast/*.bam \
 -o minimap2/yeast/transcript_ends.tsv.gz
~~~

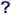 **TROUBLESHOOTING**

84. Assign each reads to a transcript using IsoQuant

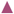 **CRITICAL** This step is not needed for less complex organisms, such as yeast. However, it needs to be performed with data from complex transcriptomes for which there are several transcripts annotated per gene, such as in zebrafish, mouse or human.

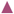 **CRITICAL** Official annotation for *S. cerevisiae* does not include UTRs (lack of TSS/TES), therefore IsoQuant will mark most of the mRNA reads as *inconsistent*.

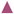 **CRITICAL** Isoquant output has to be written to a local filesystem (HDD/SDD) as it will otherwise crash if it is written to a remote filesystem such as NFS.

~~~
isoquant.py --complete_genedb --data_type nanopore --stranded reverse \
 -r ref/yeast.fa.gz -g ref/yeast.gtf.gz --bam minimap2/yeast/*.bam \
 -o isoquant
zgrep -v '^#’ isoquant/OUT/OUT.read_assignments.tsv.gz \
 | cut -f1,4,6,9 | gzip > minimap2/yeast/read_assignments.tsv.gz
~~~

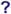 **TROUBLESHOOTING**

85. Estimate polyT tail length and composition combining all above information. 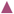 **CRITICAL** R9 run was sequenced using an old Nano3P-seq primer. Alternative Nano3P-seq primer sequence can be provided using *-p*. Resulting output can be visualized as shown in **Box 2**.

~~~
f=minimap2/yeast/R9.bam;
python ../src/get_pT.py -o $f.pT.tsv.gz -b $f \
 -e minimap2/yeast/transcript_ends.flt.tsv.gz.bed \
 -i minimap2/yeast/read_assignments.tsv.gz \
 -p CAGCACCTCTTCCGATCACTTGCCTGTCGCTCTATCTTC
f=minimap2/yeast/R10.bam;
python ../src/get_pT.py -o $f.pT.tsv.gz -b $f \
 -e minimap2/yeast/transcript_ends.flt.tsv.gz.bed \
 -i minimap2/yeast/read_assignments.tsv.gz
~~~

##### Box 2

**Visualizing PolyTailor output** 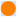 **Timing** 0.1 h

To visualize PolyTailor output, open the .tsv file in a text editor such as Vim or bash, or in Microsoft Excel.

The image below shows a part of the PolyTailor output from the yeast total RNA Nano3P-seq sequencing dataset.

More examples are available at https://github.com/novoalab/polyTailor/tree/main/test.

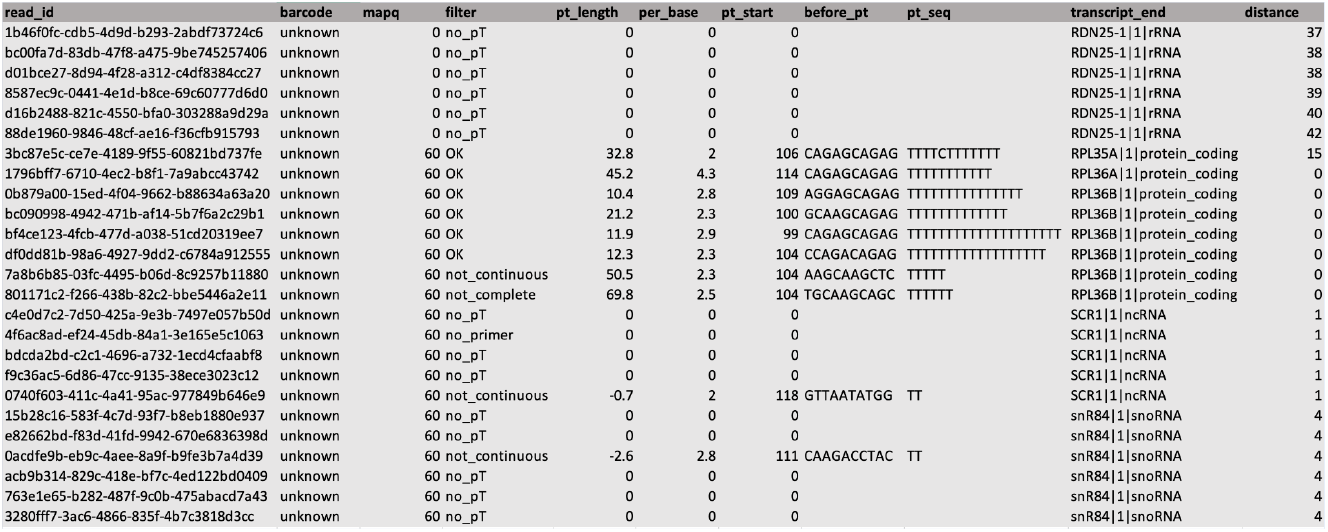

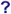 **TROUBLESHOOTING**

#### Troubleshooting

Troubleshooting guidelines can be found in **Table 1**.

**Table 1.**
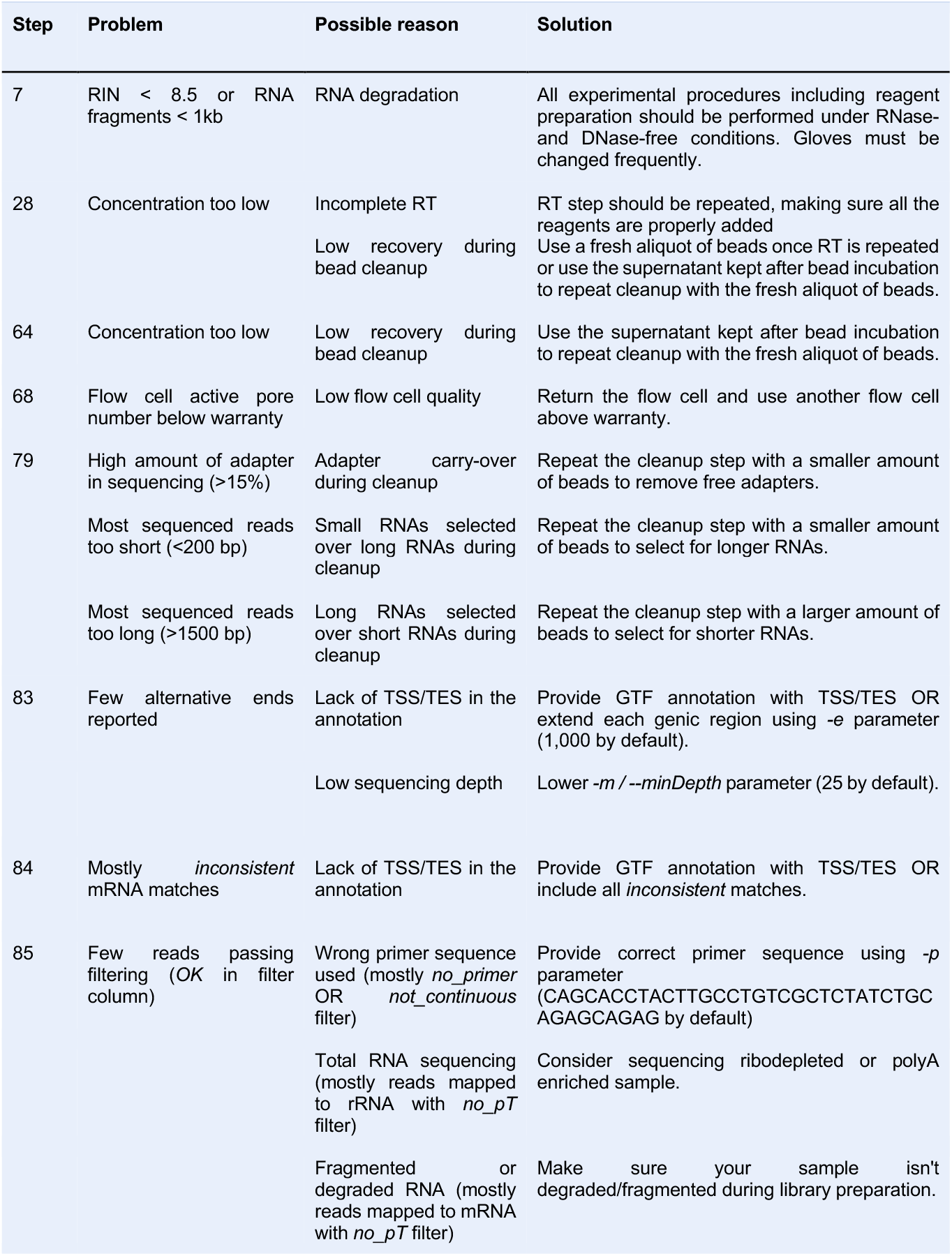
Troubleshooting table.

#### Timing

Steps 1-7, Total RNA extraction: 2 h

Steps 8-9, Pre-annealing of the oligos: 30 m

Steps 10-28, Reverse Transcription by template switching: 1 h

Steps 29-48, Complementary DNA annealing and Native Barcode Ligation: 1 h

Steps 49-64, Native Adapter (NA) Ligation: 1 h

Steps 65-79, Library loading and sequencing: 50 h

Steps 80-85, Data analysis: 2h (hac) - 15h (sup)

#### Anticipated results (<800 words)

Before moving onto a biological model, we benchmarked the new version of the Nano3P-seq protocol and of our new *in-house* software, *PolyTailor*, using DNA standards with known tail lengths and tail compositions, which were already employed to benchmark Nano3P-seq in our previous work ^8^. Each ‘DNA tail length standard’ contains a different polyT stretch length, thus allowing for assessing the tail length estimation accuracy of *PolyTailor*, whereas each ‘DNA tail content standard’ contains different tail compositions for assessing the ability of *PolyTailor* to capture tail content information **(Figure 4a)**. Our results showed that *PolyTailor* was able to accurately recapitulate polyT tail length estimates in each standard **(Figure 4b)**. Moreover, we confirmed that *dorado* basecaller did not accurately report the length of the polyT stretch, in agreement with our previous observations when using the *guppy* basecaller ^8^ **(Figure 4c)**. *PolyTailor* also accurately recapitulated the tail composition of the DNA tail content standards (**Figure 4d-e)**. Overall, our results demonstrate that our updated Nano3P-seq protocol, coupled to PolyTailor, can accurately capture tail length and content information from Nano3P-seq cDNA reads.

**Figure 4.**
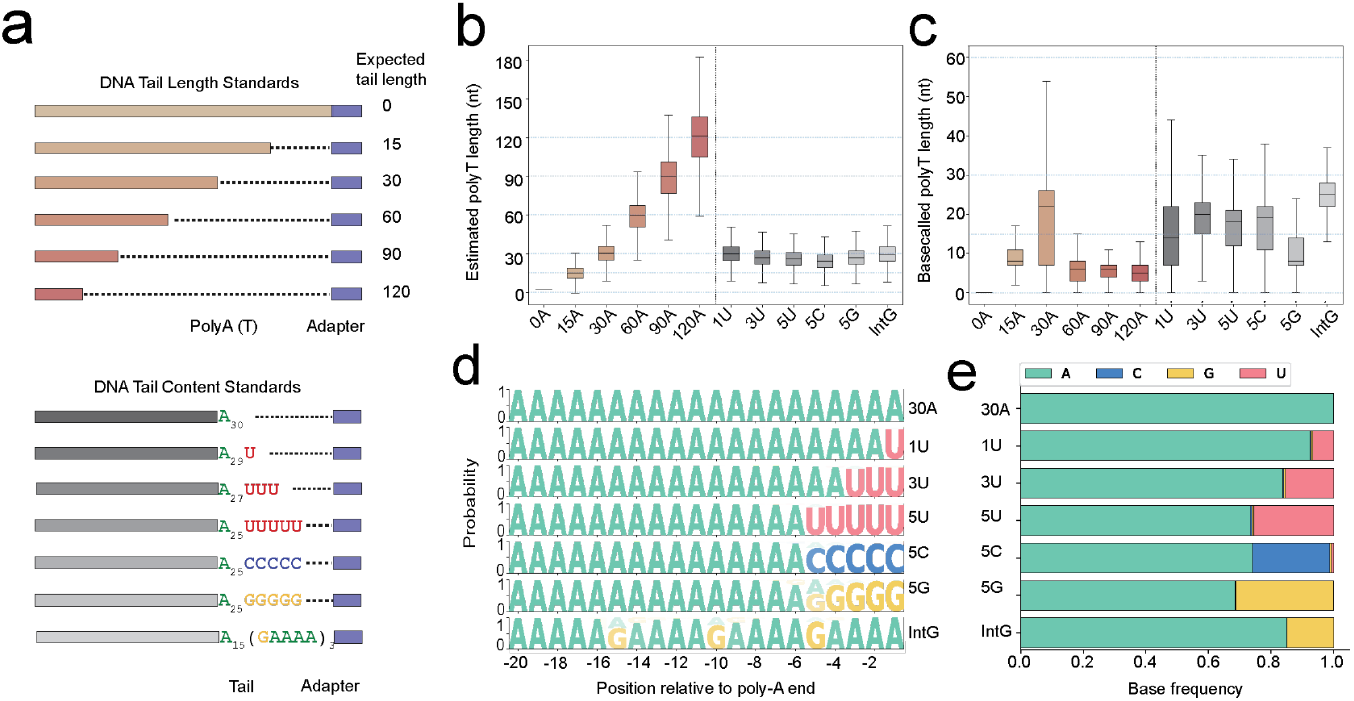
Benchmarking the new protocol and PolyTailor. **(a)** DNA tail length and content standards are synthetic DNA molecules of known varying tail lengths and tail composition. **(b)** Boxplot depicting tail length estimates (nt) reported by PolyTailor for DNA standards with known tail lengths, matching the expected results. The box extends from the first quartile to the third quartile of the data, with a line at the median. The whiskers extend from the box to the farthest data point lying within 1.5x the interquartile range from the box. **(c)** Boxplot depicting the basecalled polyT length (nt) predicted for each DNA standard, showing that basecalling cannot be used to obtain reliable estimates of poly(A) tail lengths. **(d)** Motif plot depicting the base probability for the last 20 bases of each of the DNA standards. **(e)** Base frequency bar plot illustrating the percentage of nucleotides in the tail of each DNA standard. A minimum of 10,000 reads were analyzed for each DNA tail standards in the plots included in this figure.

We then tested the Nano3P-seq protocol using total RNA from *S. cerevisiae*. We should note that isolating total RNA from yeast cultures, the expected minimum amount of total RNA in Steps 1-7 is 1 ug for flash frozen yeast cells, with a minimum RIN value of 8.5 (**Figure 5a**). In Steps 8-28, the amount of reverse transcription product should be around 5 ng from 50 ng starting material (**Figure 5b**) After Complementary DNA annealing and Native Barcode Ligation (Steps 29-48), the concentration of the ligated product is expected to be around 2-5 ng. Upon Native Adapter (NA) ligation (Steps 49-64), the concentration of the ligated product is expected to be around 2 ng or less. Library loading and sequencing steps take up to 2 days with an expected throughput of at least 5 million reads (Steps 65-79).

**Figure 5.**
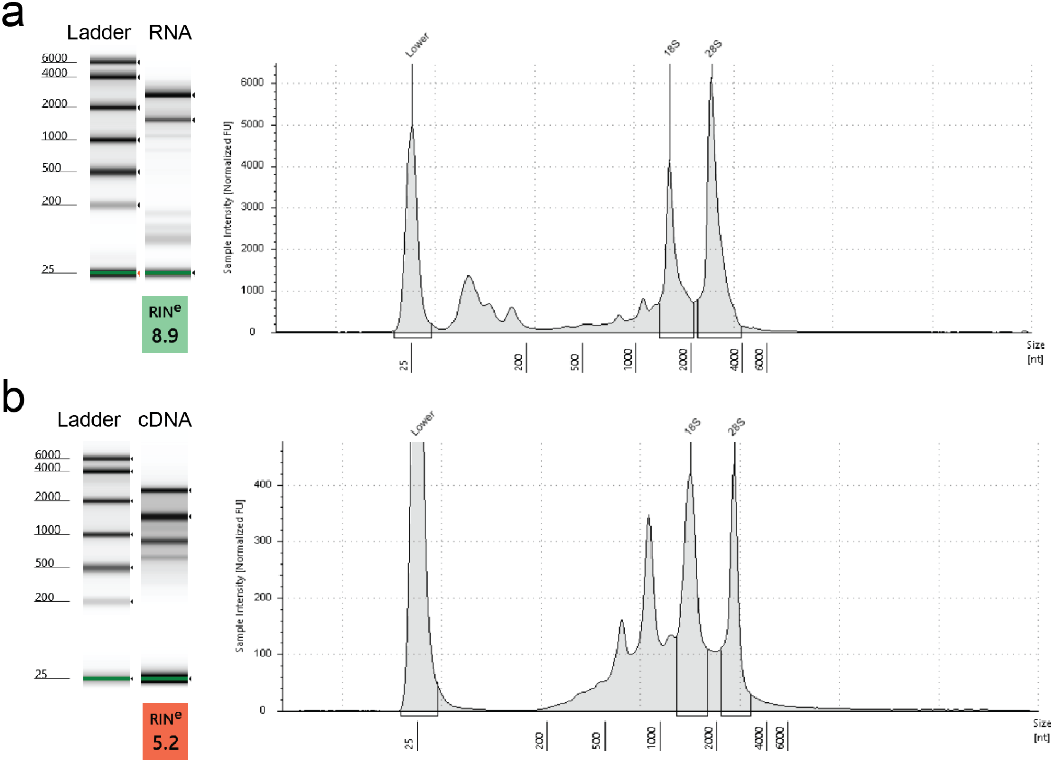
Anticipated RNA and cDNA profile. **(a)** Size distribution of extract Yeast total RNA and **(b)** cDNA synthesized from this input , both obtained using TapeStation (Agilent). *Left panels*: Electronic gel image with an electronic ladder depicts the RIN value and traditional size distribution *Right panels*: Electropherogram showing the size density distribution of the same sample.

Upon the completion of sequencing, data analysis steps include basecalling and aligning nanopore cDNA reads, classifying them and estimating their tail status (Steps 80-85). We expect Nano3P-seq to capture both non-polyA and polyA tailed RNAs from the yeast total RNA sample. Biotype fraction differences are to be expected between TGIRT and Induro RT enzymes, due to their slight differences **(Figure 6a)**. This, however, does not result in differences in mRNA count estimation, as their mRNA count profile shows high correlation (Pearson correlation r: 0.96) **(Figure 6b)**.

**Figure 6.**
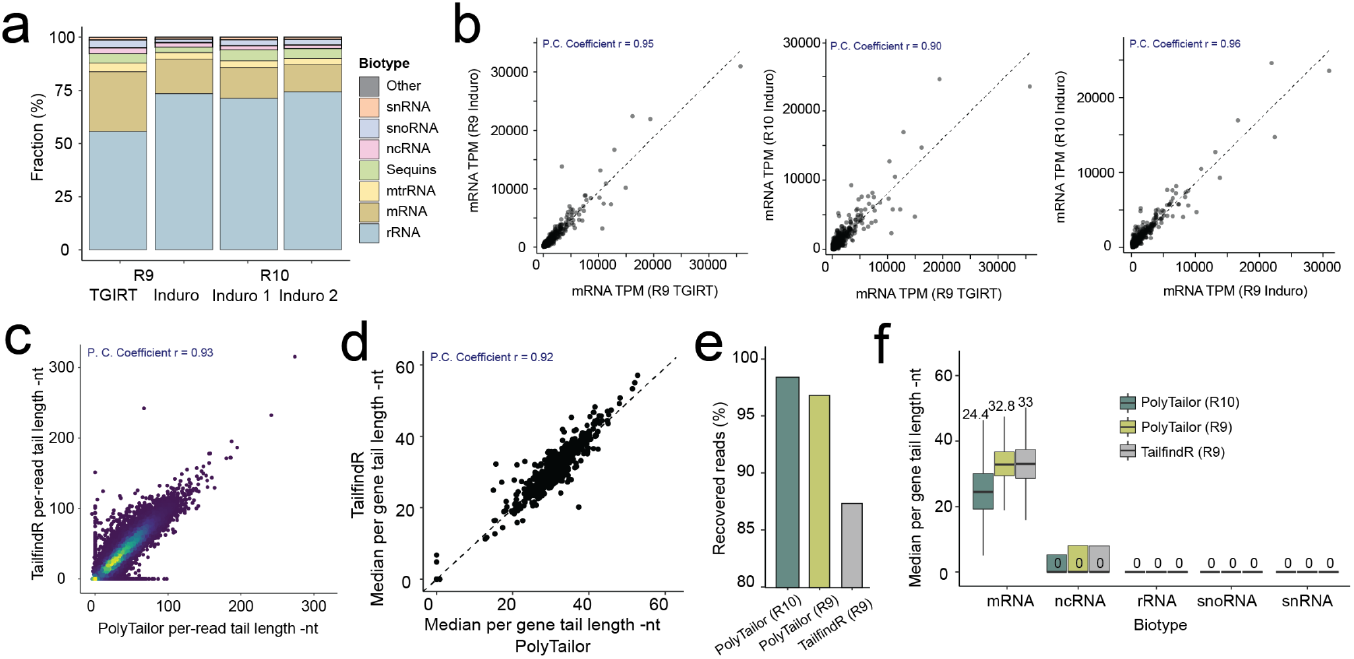
Anticipated results from data analysis. (a) Barplot showing the fraction of each biotype sequenced with Nano3P-seq protocol using TGIRT and Induro, in R9 and R10 chemistry. (b) Scatter plot illustrating the comparison between scaled counts of each yeast mRNA obtained with Nano3P-seq protocol using TGIRT and Induro, in R9 and R10 chemistry. Each dot represents an mRNA. (c) Density scatter plot illustrating the comparison between PolyTailor and tailfindR tail length estimations for the same read. (d) Scatter plot illustrating the comparison between PolyTailor and tailfindR median tail length estimates per gene. (e) Barplot showing the amount of reads recovered with a tail estimation for further analysis using PolyTailor or tailfindR. (f) Boxplot illustrating the distribution of median tail length estimates for each biotype using PolyTailor or tailfindR.

Upon *PolyTailor*, per-read predictions are expected to correlate well with tailfindR estimations used in the previous study ^8^ **(Figure 6c)**. We expect this observation to be valid for per gene median tail length estimations **(Figure 6d)**. PolyTailor tool is expected to recover more reads for which a tail can be predicted, compared to tailfindR, as well as compared to previous R9 chemistries **(Figure 6e)**. Finally, mRNA molecules are expected to contain a polyA tail, whereas RNA biotypes such as ncRNAs, rRNAs, snRNAs and snoRNAs are not expected to contain polyA tails in *S. cerevisiae* **(Figure 6f)**.

## Data availability

The data used in this study has been deposited in the European Nucleotide Archive (ENA) at EMBL-EBI under accession number PRJEB80101. A subset of the Nano3P-seq reads used in this manuscript (and their respective reference files) can be downloaded from the public repository of Novoa Lab for pipeline testing: https://public-docs.crg.es/enovoa/public/lpryszcz/src/polyTailor/test/.

## Code availability

The PolyTailor software used for Nano3P-seq analysis, including test data, has been made publicly available in GitHub at https://github.com/novoalab/polyTailor ^28^

## Acknowledgements

This work was supported by the Spanish Ministry of Science, Innovation and Universities (MCIN/AEI/10.13039/501100011033/ FEDER, UE) (PID2021-128193NB-100 to EMN). This project has received funding from the European Union’s Horizon Europe under the grant agreement No 101042103 to EMN). Funded by the European Union. Views and opinions expressed are however those of the author(s) only and do not necessarily reflect those of the European Union. Neither the European Union nor the granting authority can be held responsible for them. OB was supported by CRG Proof-Of-Concept funding. We acknowledge support of the Spanish Ministry of Science and Innovation through the Centro de Excelencia Severo Ochoa (CEX2020-001049-S, MCIN/AEI /10.13039/501100011033), the Generalitat de Catalunya through the CERCA programme and to the EMBL partnership. We are grateful to the CRG Core Technologies Programme for their support and assistance in this work.Figures for the paper were drawn using BioRender.

## Author contributions

OB adapted the experimental protocol for Nano3P-seq for R10 flowcells. LPP developed the *PolyTailor* software. OB, AMN, EV and EMN troubleshooted the tail length prediction tool development. OB built the figures. EMN supervised the project. OB, LP and EMN wrote the manuscript.

## References

1. Weill, L., Belloc, E.Bava, F.-A. & Méndez, R. Translational control by changes in poly(A) tail length: recycling mRNAs. Nat Struct Mol Biol 19, 577–585 (2012).

2. Passmore, L. A. & Coller, J. Roles of mRNA poly(A) tails in regulation of eukaryotic gene expression. Nature Reviews Molecular Cell Biology 23, 93–106 (2021).

3. Brouze, A., Krawczyk, P. S., Dziembowski, A. & Mroczek, S. Measuring the tail: Methods for poly(A) tail profiling. Wiley Interdisciplinary Reviews: RNA 14, e1737 (2023).

4. Liu, J. & Lu, F. Beyond simple tails: poly(A) tail-mediated RNA epigenetic regulation. Trends in Biochemical Sciences 49, 846–858 (2024).

5. Lucas, M. C. & Novoa, E. M. Long-read sequencing in the era of epigenomics and epitranscriptomics. Nat. Methods 20, 25–29 (2023).

6. Jain, M., Olsen, H. E., Paten, B. & Akeson, M. The Oxford Nanopore MinION: delivery of nanopore sequencing to the genomics community. Genome Biol. 17, 239 (2016).

7. Garalde, D. R. et al. Highly parallel direct RNA sequencing on an array of nanopores. Preprint at 10.1101/068809.

8. Begik, O. et al. Nano3P-seq: transcriptome-wide analysis of gene expression and tail dynamics using end-capture nanopore cDNA sequencing. Nature Methods 20, 75–85 (2022).

9. Oikonomopoulos, S. et al. Methodologies for Transcript Profiling Using Long-Read Technologies. Front. Genet. 11, 606 (2020).

10. Ramsköld, D. et al. Full-length mRNA-Seq from single-cell levels of RNA and individual circulating tumor cells. Nat. Biotechnol. 30, 777–782 (2012).

11. Au, K. F. et al. Characterization of the human ESC transcriptome by hybrid sequencing. Proc. Natl. Acad. Sci. U. S. A. 110, E4821–30 (2013).

12. Hardwick, S. A. et al. Spliced synthetic genes as internal controls in RNA sequencing experiments. Nat. Methods 13, 792–798 (2016).

13. Begik, O. et al. Nano3P-seq: transcriptome-wide analysis of gene expression and tail dynamics using end-capture nanopore sequencing. bioRxiv 2021.09.22.461331 (2021) doi:10.1101/2021.09.22.461331.

14. Legnini, I., Alles, J., Karaiskos, N., Ayoub, S. & Rajewsky, N. FLAM-seq: full-length mRNA sequencing reveals principles of poly(A) tail length control. Nat. Methods 16, 879–886 (2019).

15. Liu, Y., Nie, H., Liu, H. & Lu, F. Poly(A) inclusive RNA isoform sequencing (PAIso-seq) reveals wide-spread non-adenosine residues within RNA poly(A) tails. Nat. Commun. 10, 5292 (2019).

16. Liu, Y., Nie, H., Zhang, Y., Lu, F. & Wang, J. Comprehensive analysis of mRNA poly(A) tails by PAIso-seq2. Sci. China Life Sci. 66, 187–190 (2023).

17. Long, Y., Jia, J., Mo, W., Jin, X. & Zhai, J. FLEP-seq: simultaneous detection of RNA polymerase II position, splicing status, polyadenylation site and poly(A) tail length at genome-wide scale by single-molecule nascent RNA sequencing. Nat. Protoc. 16, 4355–4381 (2021).

18. Jia, J. et al. An atlas of plant full-length RNA reveals tissue-specific and monocots-dicots conserved regulation of poly(A) tail length. Nat. Plants 8, 1118–1126 (2022).

19. Gumińska, N. et al. LRB-IIMCB/ninetails: v.1.0.2_manuscript. (Zenodo, 2024). doi:10.5281/ZENODO.13309819.

20. Nottingham, R. M. et al. RNA-seq of human reference RNA samples using a thermostable group II intron reverse transcriptase. RNA 22, 597–613 (2016).

21. Wu, D. C. & Lambowitz, A. M. Facile single-stranded DNA sequencing of human plasma DNA via thermostable group II intron reverse transcriptase template switching. Sci. Rep. 7, 8421 (2017).

22. Xu, H., Yao, J., Wu, D. C. & Lambowitz, A. M. Improved TGIRT-seq methods for comprehensive transcriptome profiling with decreased adapter dimer formation and bias correction. Sci. Rep. 9, (2019).

23. Mohr, S. et al. Thermostable group II intron reverse transcriptase fusion proteins and their use in cDNA synthesis and next-generation RNA sequencing. RNA 19, 958–970 (2013).

24. Lentzsch, A. M., Yao, J., Russell, R. & Lambowitz, A. M. Template-switching mechanism of a group II intron-encoded reverse transcriptase and its implications for biological function and RNA-Seq. J. Biol. Chem. 294, 19764 (2019).

25. Behrens, A., Rodschinka, G. & Nedialkova, D. D. High-resolution quantitative profiling of tRNA abundance and modification status in eukaryotes by mim-tRNAseq. Mol. Cell 81, (2021).

26. Prjibelski, A. D. et al. Accurate isoform discovery with IsoQuant using long reads. Nat. Biotechnol. 41, 915– 918 (2023).

27. Xu, H., Nottingham, R. M. & Lambowitz, A. M. TGIRT-seq Protocol for the Comprehensive Profiling of Coding and Non-coding RNA Biotypes in Cellular, Extracellular Vesicle, and Plasma RNAs. Bio-protocol 11, (2021).

28. Begik, O., Pryszcz, L. P., Niazi, A. M., Valen, E. & Novoa, E. M. Nano3P-seq: charting the coding and noncoding transcriptome at single molecule resolution. Zenodo (2024) doi:10.5281/zenodo.14082765.

